# A novel mapping strategy utilizing mouse chromosome substitution strains identifies multiple epistatic interactions that regulate complex traits

**DOI:** 10.1101/2020.04.10.034637

**Authors:** Anna K. Miller, Anlu Chen, Jacquelaine Bartlett, Li Wang, Scott M. Williams, David A. Buchner

**Author notes:** Corresponding author’s contact information: David A. Buchner, Case Western Reserve University School of Medicine, 10900 Euclid Ave., Cleveland, OH, 44106-4935. These authors contributed equally (Scott M. Williams, David A. Buchner). Division of Endocrinology, Diabetes, and Metabolism, Beth Israel Deaconess Medical Center, and Harvard Medical School, Boston, MA 02115.

## Abstract

The genetic contribution of additive versus non-additive (epistatic) effects in the regulation of complex traits is unclear. While genome-wide association studies typically ignore gene-gene interactions, in part because of the lack of statistical power for detecting them, mouse chromosome substitution strains (CSSs) represent an alternate and powerful model for detecting epistasis given their limited allelic variation. Therefore, we utilized CSSs to identify and map both additive and epistatic loci that regulate a range of hematologic- and metabolism-related traits, as well as hepatic gene expression. Quantitative trait loci (QTLs) were identified using a CSS-based backcross strategy involving the segregation of variants on the A/J-derived substituted chromosomes 4 and 6 on an otherwise C57BL/6J genetic background. In the liver transcriptomes of offspring from this cross, we identified and mapped additive QTLs regulating the hepatic expression of 768 genes, and epistatic QTL pairs for 519 genes. Similarly, we identified additive QTLs for fat pad weight, platelets, and the percentage of granulocytes in blood, as well as epistatic QTL pairs controlling the percentage of lymphocytes in blood and red cell distribution width. The variance attributed to the epistatic QTL pairs was approximately equal to that of the additive QTLs; however, the SNPs in the epistatic QTL pairs that accounted for the largest variances were undetected in our single locus association analyses. These findings highlight the need to account for epistasis in association studies, and more broadly demonstrate the importance of identifying genetic interactions to understand the complete genetic architecture of complex traits.

## Introduction

The severity, presentation, and prevalence of complex traits and diseases are influenced by many genetic variants (Fu *et al*. 2013; Boyle *et al*. 2017). However, it remains unclear whether the variants work together in an additive manner or have non-linear effects on the phenotype, referred to as epistasis (Cordell 2002; Carlborg and Haley 2004; Jasnos and Korona 2007; Manolio *et al*. 2009; Bloom *et al*. 2013). Epistasis has been extensively observed in model organisms including *S. cerevisiae* (Jasnos and Korona 2007; Bloom *et al*. 2013; Forsberg *et al*. 2017), *C. elegans* (Lehner *et al*. 2006), *D. melanogaster* (Huang *et al*. 2012), and *M. musculus* (Shao *et al*. 2008; Tyler *et al*. 2017). In humans, epistasis has been more difficult to detect with standard genetic analyses. This is potentially due to low allele frequencies, limited sample sizes, complexity of interactions, insufficient effect sizes, diversity of genetic backgrounds, and methodological limitations due to the fact that in humans evidence for genetic effects is almost always statistical (Mackay 2014; Huang and Mackay 2016). However, despite these inherent difficulties, epistasis in humans has been detected in genome-wide interaction-based association studies and other methods for Crohn’s disease, Glaucoma, Behçet’s disease, multiple sclerosis, Hirschbrung disease, among others (Liu *et al*. 2011; Hu *et al*. 2013; Kirino *et al*. 2013; Hemani *et al*. 2014; Huang *et al*. 2015; Verma *et al*. 2016; GALARZA-MUNOZ *et al*. 2017; Chatterjee and Chakravarti 2019; Tyler *et al*. 2020). Assuming these examples generalize, discovering and understanding epistatic interactions will be critical for improving the predicting of phenotypes from genetic data as well as understanding of pathophysiology, thereby better guiding precision medicine-based decisions (Moore and Williams 2009; Cole *et al*. 2017).

One particularly powerful method of detecting epistasis in a mammalian system is based on studies of mouse chromosome substitution strains (CSSs) (Nadeau *et al*. 2000). The genomes of CSSs are comprised of one chromosome derived from a donor strain, with the remainder derived from a host strain such that when two CSSs are crossed together, any genetic interactions between the two non-homologous donor chromosomes can be more readily detected. This system provides an efficient model to map interacting quantitative trait loci (QTLs) on a fixed genetic background. This is in contrast to populations with genome-wide allelic variation that segregate in mapping populations such as the Diversity Outbred mouse population or typical analyses in humans (Churchill *et al*. 2012). Specifically, as it relates to epistasis, what might otherwise be a rare allelic combination in a segregating cross can be fixed in the genome of a CSS, allowing for detailed and reproducible studies of specific allelic combinations. Initial studies of CSSs derived from strains C57BL/6J and A/J, which differ with respect to metabolic disease and cancer risk, indicated that epistatic interactions were a dominant feature of many complex traits (Shao *et al*. 2008).

More recent work on CSS combinations with two substituted chromosomes highlighted the importance of epistasis in the regulation of complex traits and gene expression, and built upon previous studies by directly identifying pairs of QTLs (chromosomes) with non-additive phenotypic effects. These studies found that in mice carrying two A/J-derived chromosomes on an otherwise C57BL/6J genetic background, the A/J chromosomes frequently showed evidence of non-additive interactions (Chen *et al*. 2017). The patterns of interactions were largely in the direction of negative (or suppressing) epistasis, such that traits that were altered due to the effects of one of the substituted chromosomes were returned to baseline levels due to the combined effects of the two interacting substituted chromosomes (Chen *et al*. 2017). These findings indicated that epistatic interactions can be a common feature of complex traits as well as important for maintaining homeostasis.

A major limitation to previous analyses of epistasis using CSSs is that the resolution of QTLs mapped was at the size of the entire substituted chromosomes. However, to identify the molecular nature of such QTLs requires identification of the underlying genes, which necessitates higher resolution mapping studies. Therefore, to identify novel epistatic QTLs that regulate complex traits and gene expression levels with greater mapping resolution, we generated and analyzed the N2 generation mice from a modified backcross based on using some of the same CSSs in Chen *et al*. (2017). The resulting analyses demonstrate the ability of a CSS-based backcross to detect and fine-map epistatic QTLs for complex traits. The analyses discovered significant epistatic interactions that control lymphocyte quantity and the distribution of red cell width, and the expression of hundreds of genes in the liver. Studies of this type are essential to make progress towards the molecular characterization of epistatic interactions that will provide insight into the biological pathways of disease pathophysiology as well as improve our understanding of trait heritability and genetic architecture.

## Materials and Methods

### Mice

Strains C57BL/6J-Chr6^A/J/^NaJ (B6.A6) (stock #004384), C57BL/6J-Chr4^A/J/^NaJ (B6.A4) (stock #004382), and C57BL/6J (B6) (stock #000664) were purchased from The Jackson Laboratory. Mice were maintained by brother-sister matings with offspring weaned at 3 weeks of age. Mice were housed in ventilated racks with access to food and water *ad libitum* and maintained at 21°C on a 12-hour light/12-hour dark cycle. All mice were cared for as described under the Guide for the Care and Use of Animals, eighth edition (2011) and all experiments were approved by IACUC (protocol #2016-0064) and carried out in an AAALAC approved facility. Mice were anesthetized with isoflurane prior to retro-orbital bleeding and subsequently euthanized under anesthesia by cervical dislocation for tissue collection.

### Genotyping

Genomic DNA was isolated from mouse tail tissue using DNeasy Blood and Tissue Kits (Qiagen) following digestion with proteinase K. N2 offspring of the CSS backcross were genotyped with 57 SNP markers on Chromosome 4 and 50 SNP markers on Chromosome 6 (∼1 marker per 1.5 cM) using the PlexSeq technology (Agriplex Genomics Inc.) (Kayima *et al*. 2017; Soong *et al*. 2018). Genotyping results are listed in Supplemental Table 1. Of the 150 samples, one sample did not pass the PlexSeq quality control (<66% of genotypes called) and was not used in these analyses.

### Mouse Phenotyping

Five-week old mice were fasted overnight prior to phenotypic analysis. Mice were anesthetized with isoflurane and measurements were collected for total body weight and nose to anus length. Body mass index (BMI) was calculated as g/cm^2^. Blood was collected retro-orbitally and glucose levels were measured using an OneTouch Ultra2 meter (LifeScan, Milpitas, CA, USA). Mice were subsequently euthanized by cervical dislocation, gonadal fat pads were removed and weighed, and the caudate lobe of the liver was collected and immediately placed in RNAlater Stabilization Solution (Thermo Fisher Scientific). Complete blood counts were performed on either a Heska HemaTrue or Drew Scientific Hemavet 950 blood analyzer and included measurements of white blood count (wbc), neutrophils (ne), lymphocytes (ly), monocytes (mo), red blood count (rbc), hemoglobin (hb), hematocrit (hct), mean corpuscular volume (mcv), mean corpuscular hemoglobin (mch), mean corpuscular hemoglobin concentration (mchc), red cell distribution width (rdw), platelet count (plt), mean platelet volume (mpv), granulocyte % (gran%), granulocyte KuL (gran KuL), and red cell distribution width (absolute) (rdwa).

### Transcriptome analysis

Total RNA was isolated from liver using PureLink RNA mini kit (Thermo Fisher Scientific). RNA quality was analyzed using an Agilent TapeStation with all resulting RNA Integrity Numbers (RIN) greater than 8.5. Sequencing libraries were generated from total RNA (1 ug) using the TruSeq stranded mRNA kit (Illumina Corp.) and were sequenced using a NovaSeq6000 S4. Paired-end sequencing was performed at an average read number of 51,404,587 ± 12,403,503 reads per sample with an average of 92.7% of bases reaching quality scores of at least Q30. RNA quality control, library construction, and sequencing were performed by Macrogen Inc (now Psomagen) of Rockville, Maryland.

Prior to gene expression analysis, Fastqc (version 0.11.8) was used to determine the sequence quality of the fastq files before and after adapters were trimmed with Trimmomatic (version 0.36) (Andrews 2010; Bolger *et al*. 2014). Sequencing reads were aligned to the C57BL/6J mouse genome by Hisat2 (version 2.1.0) using the Ensembl GRCm38.p5 primary DNA assembly reference sequence (Kim *et al*. 2015; Cunningham *et al*. 2019). On average, the overall alignment rate was 98.1% ± 0.3% [range: 96.9-98.6%]. Sequence alignment files were written into bam files, sorted, and indexed by Samtools (version 1.9.0) and genomic reads were counted by HTSeq (version 0.11.1) (Anders *et al*. 2015; Morgan *et al*. 2019). Genes with genomic read counts of 0 or a low yield were identified and removed from the analysis by EdgeR (version 3.10) as low yield genes provide little evidence for differential expression, followed by normalization for the remaining expressed genes (Mccarthy *et al*. 2012).

### QTL mapping

Trait data was analyzed from offspring of the CSS backcross (n=149) (Figure 1). To generate N2 offspring for analysis, B6.A4 mice and B6.A6 mice were first separately backcrossed to B6 mice. The resulting (B6.A4 x B6)F1 and (B6.A6 x B6)F1 offspring were then intercrossed to each other (Figure 1A). Their offspring, which we refer to as the N2 generation, carried one chromosome 4 and one chromosome 6 that was entirely of B6 origin and one chromosome for each of these homologues that was recombinant, partially of B6 and partially of A/J origin (Figure 1B). The remainder of the genome, excluding chromosomes 4 and 6, is entirely from B6. Trait data was measured and analyzed for 149 N2 offspring at 5 weeks of age. All trait data for all samples can be found in Supplemental Table 2. Additionally, summary statistics, Student t-tests, and Wilcoxon tests were performed in R (version 3.5.3). Gene expression was analyzed from a subset of the backcross offspring (n=49).

**Figure 1.**
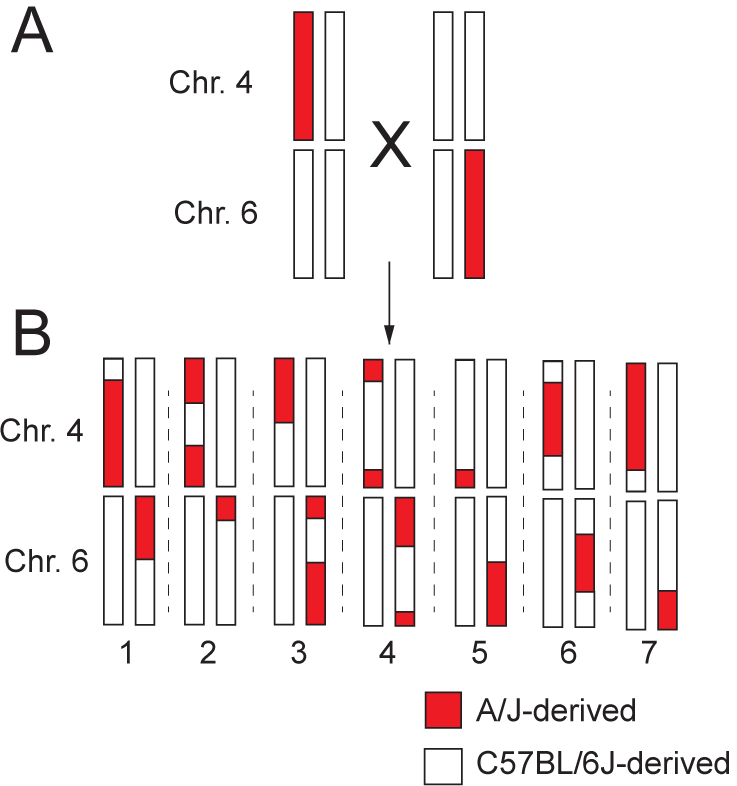
Diagram of the CSS backcross strategy to map epistatic QTL pairs. (A) (B6.A4 x B6)F1 mice were crossed with (B6 x B6.A6)F1 mice to generate 149 “N2” offspring. (B) Seven hypothetical “N2” offspring are shown. The recombinant chromosomes 4 and 6 are derived from A/J or C57BL/6J as indicated. All other chromosomes are B6-derived in all mice and therefore lack any allelic variation in this cross.

QTL linkage analysis was performed using R/qtl software (v.1.44-9), freely-available statistical software that performs single and multiple QTL mapping analyses and allows for the inclusion of covariates (Broman *et al*. 2003). One-dimensional (one locus) scans for main effects and two-dimensional (two loci) scans for epistatic effects were conducted for both the trait data and the transcriptome data to detect and map main and epistatic effects using the R/qtl functions “scanone” and “scantwo”, respectively. As some metabolism- and blood-related traits vary by sex, the scans were performed with and without sex as a covariate using logistic regression with the HK algorithm (one-dimensional) or the EM algorithm (two-dimensional). Two-dimensional scans report the interaction with the largest LOD score for each possible chromosomal interaction: Chromosome 4 to Chromosome 4, Chromosome 6 to Chromosome 6, and Chromosome 4 to Chromosome 6. Trait data was also performed with BMI as an additive covariate to account for variation due to adiposity (SAMOCHA-BONET *et al*. 2008; Marginean *et al*. 2019). LOD significance thresholds were obtained from permutation tests for both additive and epistatic models. The R/qtl permutation test permutes the phenotypes relative to the genotype data, applies the QTL mapping method to the shuffled version of the data to obtain a set of LOD curves, and derives the genome-wide maximum LOD score. Permutation tests for each trait were performed with 10,000 permutations; thus, concerns of multiple testing related to the number of statistical tests performed for each trait are accounted for by using permutation p-values (Broman 2009). QTL locations were updated in the context of a multiple QTL model based on maximum likelihood using an iterative scan with the “refineqtl” function in R/qtl (Broman *et al*. 2019). Intra-chromosomal interacting QTLs regulating gene expression within 30 MB of each other were removed as visual inspection of the data suggested a high frequency of suspected false positives, which is also consistent with the limited mapping resolution given the study’s sample size. Significant interactions were visualized in three-dimensional space using the R packages “circularize” (Gu 2014) and “rgl” (Adler 2019).

### Data availability

All RNA sequencing files can be found at the NCBI gene expression omnibus. All genotype data can be found in Supplemental Table 1 and all phenotype data can be found in Supplemental Table 2. The complete QTL analysis pipeline can be found on protocols.io.

## Results

### Identification of additive QTLs for complex traits

B6.A6 and B6.A4 mice have previously been found to carry allelic variation on their respective substituted chromosomes that non-linearly interacted with each other to regulate plasma glucose levels. This particular strain combination demonstrated the third most statistically significant interaction regulating glucose levels among 15 strain combinations tested, with evidence of this interaction found in both male and female mice. Additionally, mRNA expression of 83 genes were identified that were regulated by epistatic interactions between these two CSSs (Chen *et al*. 2017). Beyond the epistatic interactions regulating glucose and gene expression, previous studies also found evidence of QTLs mapped to these chromosomes for other complex traits, including a number of hematology-related traits that can quickly, quantitatively, and cost-effectively be measured in hundreds of mice (LAKE J. *et al*.). Studies of hematology-related traits in the B6.A6 strain in particular identified as many or more QTLs mapped to that substituted chromosome (Chr. 6) than were found on any other chromosome. Thus, given the previous findings of epistasis between allelic variation on Chromosomes 4 and 6, as well as the evidence for additional QTLs regulating clinically relevant traits that could be efficiently studied in a large mapping population, this particular strain combination was chosen for additional studies. To further map the genetic loci that control plasma glucose levels and other traits, a total of 17 metabolism- and blood-related traits were analyzed in N2 offspring from a modified backcross between B6.A6 and B6.A4 (Figure 1).

A single-locus QTL genome scan was performed to test for main (additive) effects in the 149 N2 mice (Broman *et al*. 2003). Unadjusted main effect QTL analysis identified significant QTLs for two traits: platelet count (plt) (LOD score > 2.04, p-value < 0.05) and granulocyte % (gran%) (LOD score > 2.02, p-value < 0.05). To test whether including sex as a covariate might affect QTL detection, we first tested for phenotypic differences in metabolism- and blood-related traits between male and female mice from the same crosses. There was a significant difference between sexes for 8 of the 17 factors tested: fasted body weight, fasting glucose, length, fat pad weight, mcv, mchc, plt, and rdwa (p < 0.05, Table S3). Thus, sex was included as a covariate to better account for sex-specific trait differences. Sex-adjusted main effect QTLs were identified for three traits: plt and gran% as before, plus an additional QTL for fat pad weight (Table 1). The 1-LOD support QTL intervals for these traits were located between 38.9 - 104.8 Mb on Chr. 6 (fat pad weight), 83.2 - 128.8 Mb on Chr. 4 (plt), and 125.0 - 146.5 Mb on Chr. 6 (gran%) (Table 1). These three loci accounted for 28.3% (fat pad weight), 29.4% (plt), and 29.0% (gran %) of their respective trait variances.

**TABLE 1.** 1-LOD SUPPORT INTERVALS FOR MAIN EFFECT TRAIT QTLS

|  | Chromosome | Marker | Position (Mb) | LOD | p-value | BMI LOD | BMI p-value |
| --- | --- | --- | --- | --- | --- | --- | --- |
| Fat Pad Weight | 6 | rs13478720 | 38.85 | 1.17 |  | 1.19 |  |
|  | **6** | **rs13478811** | **70.31** | **2.18** | **0.030** | **2.24** | **0.030** |
|  | 6 | rs13478947 | 104.84 | 1.06 |  | 1.08 |  |
| Platelets | 4 | rs13477808 | 83.18 | 1.56 |  | 1.55 |  |
|  | **4** | **rs13477911** | **111.29** | **2.61** | **0.018** | **2.64** | **0.010** |
|  | 4 | rs13477973 | 128.75 | 1.41 |  | 1.36 |  |
| Granulocyte (%) | 6 | rs3023094 | 125.04 | 1.61 |  | 1.49 |  |
|  | **6** | **rs13479068** | **137.48** | **2.64** | **0.006** | **2.57** | **0.02** |
|  | 6 | rs3090690 | 146.46 | 1.55 |  | 1.55 |  |

\*Peak SNPs are in bold with the SNPs defining the 1-LOD support interval shown above and below the peak SNP. The LOD and P-value columns include sex in the model, The BMI LOD and BMI p-value columns include sex and BMI in the model

Main effect QTL analyses were also performed with BMI in the model to account for variation due to adiposity. QTL analyses that included sex and BMI resulted in the same significant main effect QTLs as sex alone. However, there was an increase in significance for the plt QTL when BMI was added to the model (sex p value = 0.018 vs. sex and BMI p value=0.010) (Table 1), which aligns with the previously seen association between platelet count and BMI (SAMOCHA-BONET *et al*. 2008; Marginean *et al*. 2019).

### Identification of epistatic QTLs for complex traits

To test for epistatic QTL interactions, a two-dimensional (two-locus) genome scan with sex as a covariate was performed for the metabolism- and blood-related trait data from 149 N2 mice. Epistatic QTL pairs were identified for two of the traits: lymphocytes (ly%) and red cell distribution width (rdw). An inter-chromosomal interaction was discovered between loci on chromosomes 4 and 6 that regulated ly% and an intra-chromosomal interaction between two distinct loci on chromosome 6 regulated rdw (Table 2). The 1-LOD support QTL intervals for ly% were between 5.09 – 104.51 Mb on Chr. 4 and between 38.85 – 93.25 Mb on Chr. 6. The 1-LOD support interval for ly% on Chr. 4 contains a second peak (Figure 2D), however the direction of the effect on lymphocyte levels is the same for both regions, and given the overlap between the 1-LOD support intervals the possibility remains that these seemingly distinct peaks represent one signal. The 1-LOD support QTL intervals for rdw were between 3.49 – 16.06 Mb and 14.36 Mb - 17.72 Mb on Chr. 6 (Table 2).

**TABLE 2.** 1-LOD SUPPORT INTERVALS FOR EPISTATIC TRAIT QTLS

|  | Chromosome | Marker (nearest SNP) | Position (Mb) | LOD | p-value |
| --- | --- | --- | --- | --- | --- |
| Lymphocytes (%) |  |  |  |  | **0.032** |
| Interval 1 |  |  |  |  |  |
|  | 4 | rs13477536 | 5.09 | 2.86 |  |
|  | **4** | **rs13477644** | **35.28** | **4.23** |  |
|  | 4 | rs3695339 | 104.51 | 2.72 |  |
| Interval 2 |  |  |  |  |  |
|  | 6 | rs13478720 | 38.85 | 2.29 |  |
|  | **6** | **rs13478739** | **47.03** | **3.85** |  |
|  | 6 | rs13478897 | 93.25 | 1.95 |  |
| Red Cell Distribution Width |  |  |  |  | **0.036** |
| Interval 1 |  |  |  |  |  |
|  | 6 | rs13459096 | 3.49 | 3.10 |  |
|  | **6** | **rs13478633** | **12.25** | **5.69** |  |
|  | 6 | rs13478641 | 16.06 | 0.90 |  |
| Interval 2 |  |  |  |  |  |
|  | 6 | rs13478638 | 14.36 | 3.70 |  |
|  | **6** | **rs13478641** | **16.06** | **5.95** |  |
|  | 6 | rs3023064 | 17.72 | 3.91 |  |

\*Peak SNPs are in bold with the SNPs defining the 1-LOD support interval shown above and below both peak SNPs

**Figure 2.**
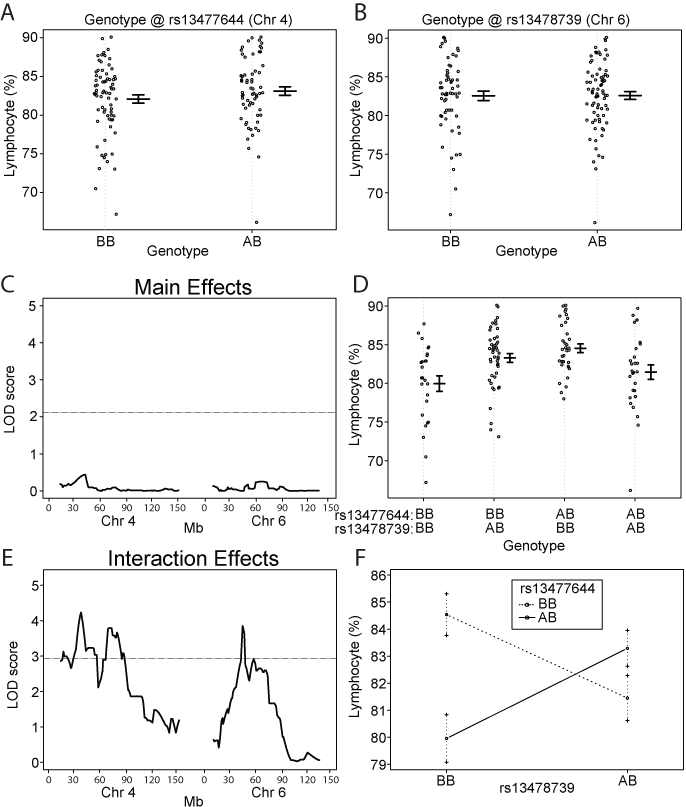
Epistatic interaction between loci on chromosomes 4 and 6 regulate the percentage of lymphocyte cells in blood. Lymphocyte % based on genotype at SNP markers (A) rs13477644 on chromosome 4 and (B) rs13478739 on chromosome 6. Each dot represents one mouse. (C) QTL mapping results for main effects on chromosomes 4 and 6. The LOD threshold for significance (p < 0.05) was calculated by permutation testing (n=10,000) and is indicated by a horizontal line. Mb position is indicated for both chromosomes 4 and 6 along the x-axis. No significant main effect QTLs were detected. (D) Lymphocyte % based on the combined genotypes at rs13477644 on chromosome 4 and rs13478739 on chromosome 6. (E) QTL mapping results for interaction effects on chromosomes 4 and 6. The LOD threshold for significance (p < 0.05) was calculated by permutation testing (n=10,000) and is indicated by a horizontal line. The most significant interaction QTL pairs were detected with peaks centered at rs13477644 (17.3 Mb) and rs13478739 (22.6 Mb), with a potentially second peak on chromosome 4 centered at rs13477796 (37.6 Mb). Mb position is indicated for both chromosomes 4 and 6 along the x-axis. (F) Context-dependent effects of the BB and AB genotypes at markers rs13477644 and rs13478739. Mean and standard error are shown for each genotype combination. An “A” genotype indicates A/J-derived. A “B” genotype indicates C57BL/6J-derived.

For ly%, the most significant interacting SNPs within the QTL intervals were rs13477644 (Chr. 4) and rs13478739 (Chr. 6). Testing for effects of these SNP genotypes individually on ly% revealed no marginal effect on lymphocyte count (Figure 2A, B). Accordingly, a main effect QTL analysis of these regions individually found no evidence for a QTL at either locus (Figures 2C, S4). However, when accounting for both SNP genotypes taken together, association with lymphocyte count became readily apparent (Figure 2D), with the resulting QTL analysis for interacting loci revealing a strong epistatic QTL (Figure 2E). The epistatic regulation of ly% represents a case of negative epistasis, as each individual A/J-derived allele at rs13477644 and rs13478739 is associated with a modest increase in ly% (Figure 2E, middle two sections relative to the left section); however, the combination of A/J-alleles at both rs13477644 and rs13478739 (Figure 2E, far right section) together reduce ly% back towards the control levels. The context dependent effects of this interaction are graphically illustrated in Figure 2F, as shown by the crossing lines connecting the mean values for each genotype combination indicating suppression epistasis. Remarkably, this epistatic interaction alone accounted for 12.1% of the variance in this trait.

For rdw, the most significant interacting SNPs within the QTL intervals were rs13478633 (Chr. 6) and rs13478641 (Chr. 6). Testing for effects of these SNP genotypes individually on rdw revealed no marginal effect on red cell distribution width (Figure S1A, B) and a main effect QTL analysis found no evidence for a QTL at either locus (Figures S1C, S4). However, when both SNP genotypes were jointly analyzed (Figure S1D), the association with red cell distribution width was identified (Figure S1E). The epistatic regulation of rdw again represents a case of negative epistasis, as each A/J-derived allele at rs13478633 and rs13478641 is associated with a modest decrease in rdw (Figure S1D, middle two sections relative to the left section); however, the combination of A/J-alleles at both rs13478633 and rs13478641 (Fig. S1D far right section) revert rdw back towards the control levels. The context dependent effects of this interaction are graphically illustrated in Figure S1F. This epistatic interaction accounted for 10.7% of variance in this trait.

### Identification of QTLs that regulate hepatic gene expression

In addition to the metabolism and blood traits, we measured the liver transcriptomes in a subset (n=49) of the offspring from the B6.A4 and B6.A6 backcross to identify additive and epistatic QTLs regulating hepatic gene expression. Linkage analyses of the RNA-Seq data revealed 768 main effect QTLs (single locus) (Table S5). Among the main effect QTLs, there were 159 genes within the QTL interval on either chromosome 4 or 6, and are thus cis-QTLs, whereas 609 were located elsewhere in the genome and are therefore trans-QTLs. There were 519 epistatic QTL pairs regulating hepatic gene expression (LOD Score > 2.44, p-value < 0.05) (Table S6). As the R/qtl mapping software used only identifies the most significant inter-chromosomal epistatic interaction between chromosome 4 and 6 and the most significant intra-chromosomal epistatic interactions for each chromosome, if anything these numbers represent an underestimate of the true number of epistatic QTL pairs, thus demonstrating the pervasive effects of epistasis in the regulation of gene expression. In addition to the detection of widespread epistasis, the N2 cross strategy used in this study (Figure 1) enabled higher resolution QTL mapping relative to prior CSS studies in which the mapping resolution was limited to entire chromosomes (Chen *et al*. 2017). For example, previous studies mapped main effect QTLs regulating the expression of *Extl1* to Chromosome 4, *Cadps2* to Chromosome 6, and an epistatic interaction controlling the expression of *Tmem245* to Chromosomes 4 and 6, all of which were mapped with higher resolution with the N2 cross (Figures S2 and S3).

The relationship between main effect QTL LOD scores and interaction QTL LOD scores shows that a large interaction QTL LOD score does not predict a large main effect QTL LOD score, and *vice versa* (Figure 3). Of the 748 unique genes with significant main effects and 510 unique genes with significant epistatic effects, 727 genes had only main effects, 489 had only epistatic effects, respectively, and 21 genes had both significant main and epistatic effects. Thus, there appears to be little overlap between marginal and epistatic effector loci (Figure 3). The percent variance explained by interaction QTLs (mean 24.5%; range 20.5% - 85.6%) was similar to that explained by single locus QTLs (mean 28.5%; range 20.5% - 44.4%). The detection of so many epistatic QTLs accounting for such a high proportion of the variance was not unexpected, as our previous data suggested the widespread importance of epistasis in regulating gene expression (Chen *et al*. 2017). Of 519 epistatic QTLs (the 510 unique genes and genes regulated by multiple eQTLs), 243 (46.8%) exhibited positive epistasis whereas 276 (53.2%) exhibited negative epistasis. That a slight majority of QTLs exhibited negative epistasis indicates that epistasis can have a profound effect on maintaining homeostasis, although the proportion of negative epistatic QTL pairs in this study was not as high as previously detected using CSSs (>95%) (Chen *et al*. 2017), However, the current proportion of negative epistasis is more in line with other studies comparing negative and positive interactions (Segre *et al*. 2005; Costanzo *et al*. 2016; Guerrero *et al*. 2017).

**Figure 3.**
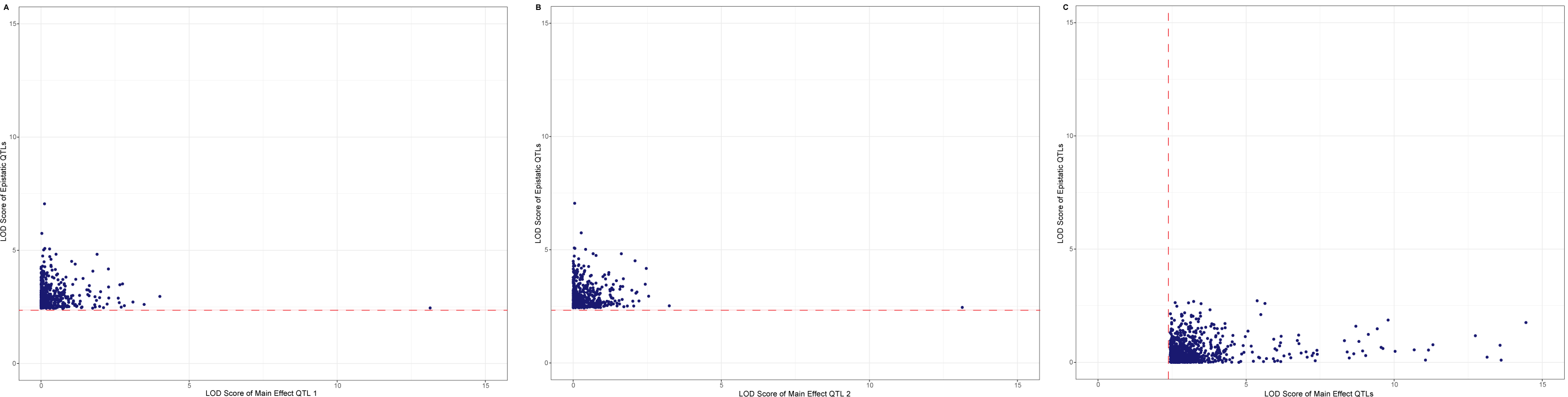
Little overlap between main effect QTLs and epistatic QTL pairs. (A,B) LOD scores for peak SNPs defining significant epistatic QTL pairs were plotted against the corresponding main effect LOD scores for each SNP within the epistatic pair. (C) LOD scores for peak SNPs defining significant main effect QTLs were plotted against the most significant interaction LOD score for that SNP. The solid horizontal red line in panels A and B indicates the threshold level for significance for epistatic QTL pairs and the vertical dashed red line in panel C indicates the threshold level for significance for main effect QTLs.

The most statistically significant epistatic QTL regulating gene expression was for *Arhgap25*, with LOD peaks at the SNPs rs13478003 (Chr. 4) and rs13478976 (Chr. 6). As seen in the analysis of ly%, analysis of *Arhgap25* expression associated with each individual SNP failed to reveal a main effect of genotype (Figure 4A, B), and neither resulted in a significant QTL association (Figure 4C). However, when *Arhgap25* expression was analyzed with SNP-SNP interactions, a clear genetic effect of the combination of rs13478003 and rs13478976 was detected (Figure 4D-F). This interaction accounted for a remarkable 41.7% of the variance for this trait, but was completely absent in a standard single locus association analysis (Table S6). Beyond *Arhgap25*, 518 additional genes were identified with significant epistatic QTLs. Interaction plots of the next 5 most significant regulated genes (after *Arhgap25*) are shown in Figure 5A-E and include *Irf2bpl, Gnmt, Rasa3, Flnb*, and *Capn10*, all demonstrating interactions that act to maintain homeostatic levels of gene expression (negative epistasis), rather than exacerbate strain differences in expression levels (Figure 5). None of the interacting loci that associated with the regulation of these transcripts was marginally associated with controlling gene expression levels.

**Figure 4.**
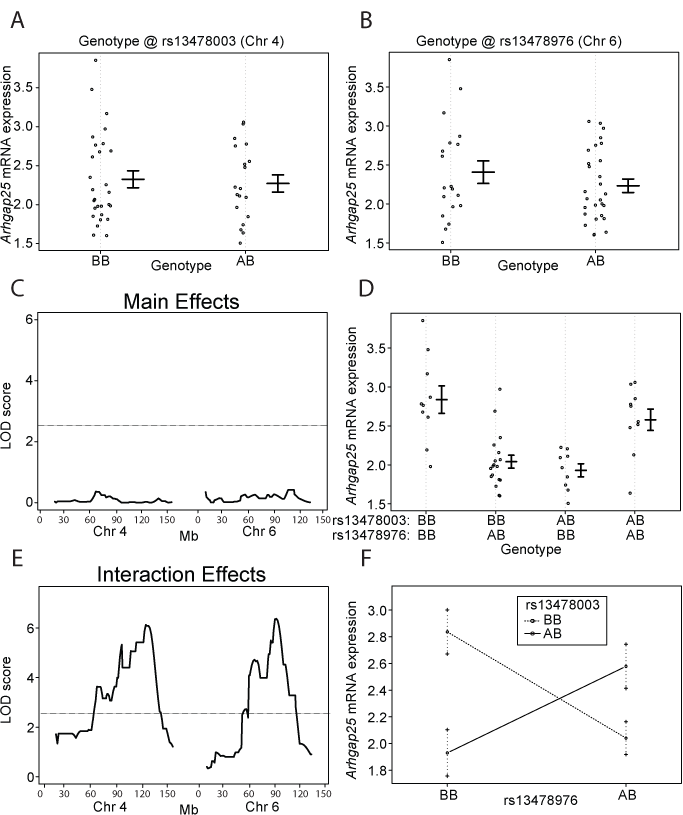
Epistatic interaction between loci on chromosomes 4 and 6 regulates *Arhgap25* mRNA expression. *Arhgap25* mRNA expression based on genotype at SNP markers (A) rs13478003 on chromosome 4 and (B) rs13478976 on chromosome 6. Each dot represents one mouse. (C) QTL mapping results for main effects on chromosomes 4 and 6. The LOD threshold for significance (p < 0.05) was calculated using R/qtl and is indicated by a horizontal line. Mb position is indicated for both chromosomes 4 and 6 along the x-axis. No significant QTLs were detected. (D) *Arhgap25* mRNA expression based on the combined genotypes at rs13478003 on chromosome 4 and rs13478976 on chromosome 6. (E) QTL mapping results for interaction effects on chromosomes 4 and 6. The LOD threshold for significance (p < 0.05) was calculated by R/qtl using 10,000 permutations and is indicated by a horizontal line. The most significant interaction QTLs were detected with peaks at rs13478003 (69.1 Mb) and rs13478976 (52.2 Mb). Mb position is indicated for both chromosomes 4 and 6 along the x-axis. (F) Context-dependent effects of the BB and AB genotypes at rs13478003 and rs13478976. Mean and standard error are shown for each genotype combination. An “A” genotype indicates A/J-derived. A “B” genotype indicates C57BL/6J-derived.

**Figure 5.**
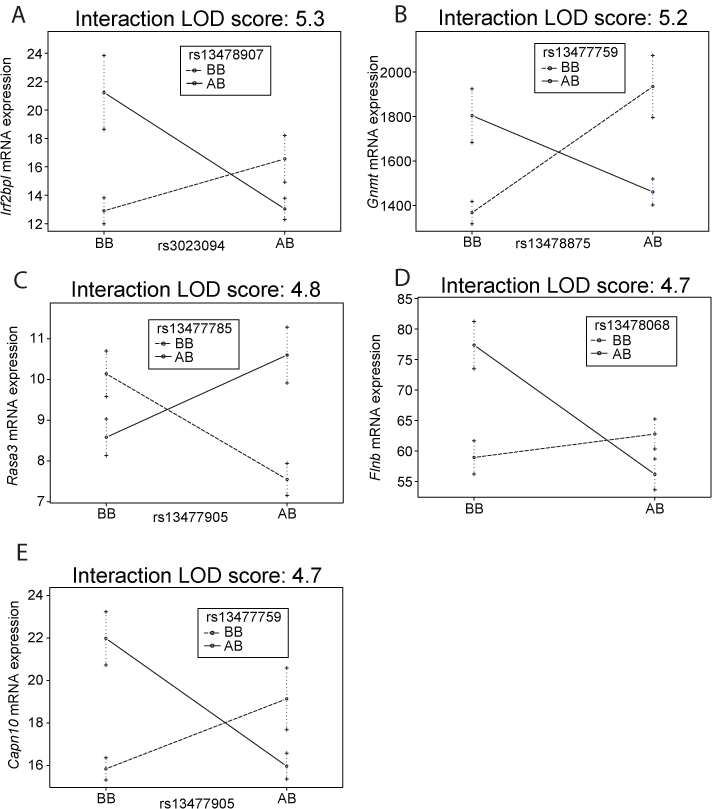
Epistatic interactions control gene expression. Context dependent effects on gene expression for (A) *Irf2bpl*, (B) *Gnmt*, (C) *Rasa3*, (D) *Flnb*, and (E) *Capn10*. The y-axis represents the number of unique sequencing reads per gene after normalizing read depth across samples. Mean and standard error are shown for each genotype combination. An “A” genotype indicates A/J-derived. A “B” genotype indicates C57BL/6J-derived.

Interacting loci on chromosomes 4 and 6 were widely distributed along each chromosome (Figure 6A). Of the 186 intra-chromosomal interactions, 37 were located on Chromosome 4 and 149 on Chromosome 6 (Figure 6B). The remaining 333 interactions were inter-chromosomal (Figure 6C). Of the genes whose expression was regulated by an epistatic QTL pair, 19 were located within the QTL interval on either chromosome 4 or 6, and are thus likely controlled by cis-QTLs. This represents a 2.0-fold enrichment of cis-QTLs based on the size of the QTL intervals and relative to the number of cis-QTLs expected by chance (p = 0.006). The remaining 500 interacting eQTLs were located elsewhere in the genome and are therefore trans-QTLs. Although there appeared to be few if any QTL hotspots, the genes that were regulated by epistatic QTL pairs were enriched for a number of biological pathways (Supplemental Table 7). Gene ontology analysis discovered that the most significantly enriched pathways tended to be non-specific pathways related to various metabolic processes. Other pathways identified were related to basic cellular processes including the regulation of gene expression and signal transduction, RNA processing, and stress responses. All this is to say that the genes regulated by epistasis collectively have broad cellular functions and represent the most fundamental cellular biological processes.

**Figure 6.**
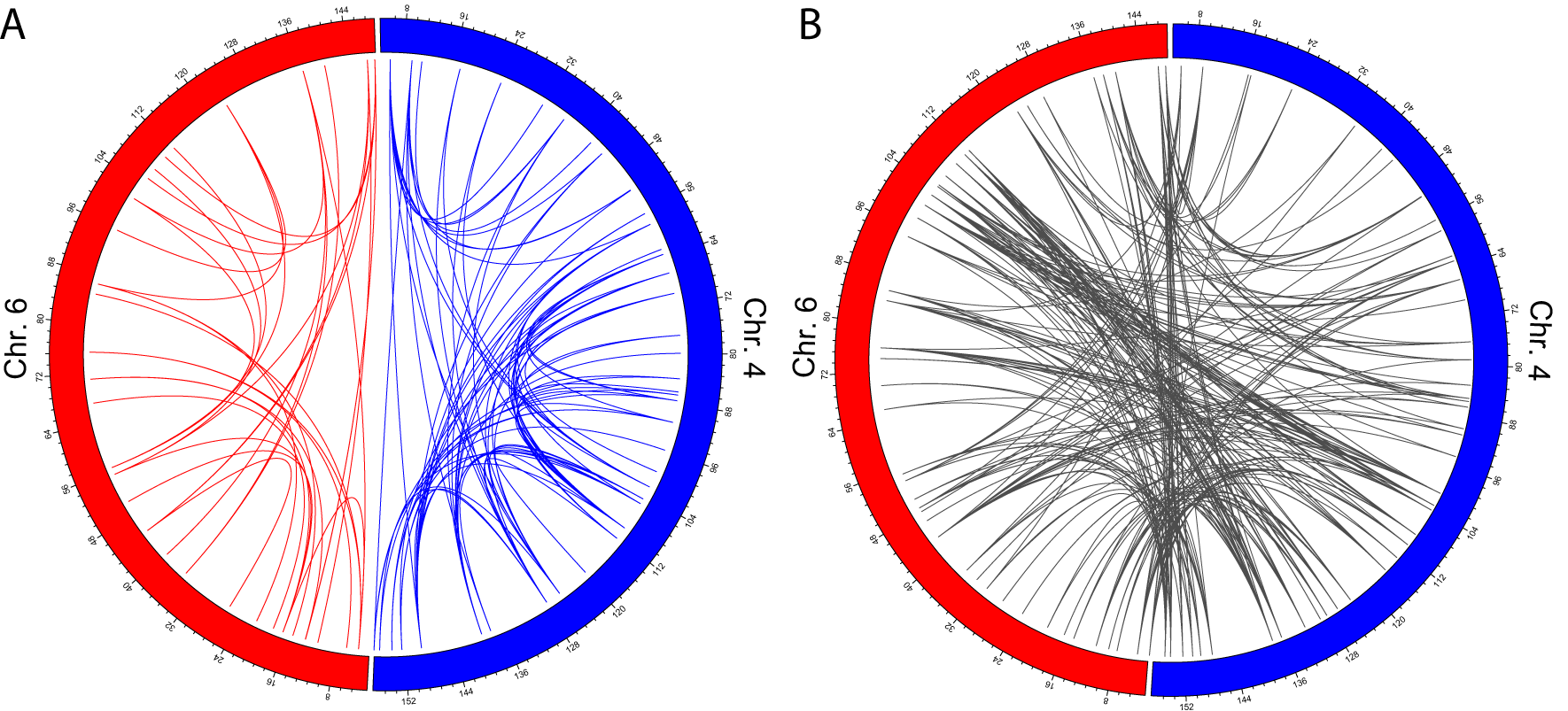
Inter-chromosomal and intra-chromosomal epistatic interactions that control gene expression are widely distributed along chromosomes 4 and 6. (A) The circos plots illustrates the location along chromosomes 4 and 6 for the peak SNPs for all epistatic QTL pairs regulating gene expression. Chromosome 6 and the intra-chromosomal epistatic QTL pairs on that chromosome (n=37) are shown in red. Chromosome 4 and the intra-chromosomal epistatic QTL pairs on that chromosome (n=149) are shown in blue. Numbering around the circle plots indicates the Mb position on each chromosome. (B) Inter-chromosomal epistatic QTL pairs between chromosomes 4 and 6 (n=333) are shown in grey.

## Discussion

N2 CSS mice with allelic variation from two substituted chromosomes were used to study epistatic interactions and map loci controlling complex traits and gene expression, as these CSSs have a greatly simplified genetic background relative to other commonly used mapping resources in mammalian systems. The resulting analyses identified widespread epistatic interactions, including specific interacting loci that contributed to the heritable variation in lymphocyte percentage (ly%), red cell distribution width (rdw), and hepatic gene expression of over 500 genes. Intra-chromosomal interactions were detected in rdw and hepatic gene expression data that could not be identified in studies of the B6.A4 and B6.A6 CSS strains carrying the entire nonrecombinant A/J-derived substituted chromosomes (Chen *et al*. 2017). That this number of traits and genes were identified with detectable regulation by epistasis is remarkable considering that only a single time point was examined, gene expression was only measured in the liver, and only one pairwise strain combination of CSSs was examined. Given that ly% and rdw are associated with diagnosis and prognosis in cancer patients as well as cardiometabolic disease, identifying a more complete genetic model that incorporates interaction may improve our ability to predict those at greatest risk or modulate these traits based on improved knowledge of the biological pathways that underlie these interactions (Iseki *et al*. 2017; Zhao *et al*. 2017; Pilling *et al*. 2018).

Among the most significant genes whose expression is regulated by epistatic interactions, *Capn10* is particularly interesting given its controversial association with T2D risk. It was the first T2D susceptibility gene identified in linkage studies (Horikawa *et al*. 2000), with multiple subsequent GWAS or candidate gene studies having also identified an association between variants near *CAPN10* and T2D risk and other diabetes-related phenotypes (Baier *et al*. 2000; Evans *et al*. 2001; Orozco *et al*. 2014; Ibrahim *et al*. 2015; Cui *et al*. 2016; Zhao *et al*. 2016; Bayramci *et al*. 2017; Hou *et al*. 2017; CASTRO-MARTINEZ *et al*. 2018). However, many other studies failed to replicate these findings (Hegele *et al*. 2001; Tsai *et al*. 2001; Khan *et al*. 2014; AL-SINANI *et al*. 2015; Plengvidhya *et al*. 2015; Zhang *et al*. 2019). It is tempting to speculate that one potential reason for the inconsistent results could be an incomplete understanding of epistatic interactions between *CAPN10* and other loci. To illustrate how this could happen, previous simulation studies of interacting loci revealed that small differences in allelic frequencies between populations can lead to a failure to replicate GWAS results (Greene *et al*. 2009). Therefore, it is of note that one study of ∼1,400 T2D and control individuals found no evidence of main effects in *CAPN10*, but did identify evidence for gene-gene interactions involving *CAPN10* influencing T2D risk (Uma JYOTHI AND REDDY 2015). This is remarkably similar to what we have shown with ly% (Figure 3) and *Arhgap25* expression (Figure 4), where a failure to account for interactions can mask an otherwise strong association.

The ∼500 genes whose hepatic expression was regulated by epistasis is significantly more than were first identified in studies of the B6.A4 and B6.A6 CSS strains carrying the entire nonrecombinant A/J-derived substituted chromosomes (Chen *et al*. 2017). This may be, in part, because the recombination events within chromosomes 4 and 6 decoupled intra-chromosomal interactions in order to reveal previously undetected intra- and inter-chromosomal epistatic QTLs. This phenomenon has been frequently observed in previous studies of CSS for a wide range of traits, with numerous previously hidden QTLs revealed upon deconstruction of the substituted chromosome (Buchner and Nadeau 2015). The high prevalence of epistatic interactions provides evidence for complex molecular models underlying the genetics of many quantitative traits (Gerke *et al*. 2009; Gerke *et al*. 2010; Chow 2016; Sackton and Hartl 2016). In addition, it is of note that there is little overlap between the genes that affect gene expression via marginal versus epistatic actions. This means that conditioning tests for epistasis based on prior detected marginal effects, as is often done, may underestimate the importance of epistasis in complex traits (Greene *et al*. 2009).

Beyond the identification of specific epistatic QTLs, this study has broad implications for understanding the genetic architecture of complex traits. For instance, among the traits where an epistatic QTL was detected, those QTLs accounted for a considerable proportion of the variance for that trait (11.4% for blood-related traits, 24.5% for gene expression). The gene expression QTLs revealed a similar number of loci with comparable variance explained (interaction: mean 24.5%; range 20.5% - 85.6%, marginal: mean 28.5%; range 20.5% - 44.4%). This demonstrates that epistatic interactions can have large effect sizes. However, it is important to note that studies in CSSs have consistently found QTLs that explain larger proportions of the variance relative to other mapping populations such as the hybrid mouse diversity panel and the diversity outbred mapping populations (Buchner and Nadeau 2015). The high variance accounted for by QTLs in studies based on CSSs has been attributed in part to the limited allelic variation within each CSS, the limited power for detecting smaller effects given study sample sizes, and the study of genetically identical replicates within each CSS strain to improve genotype-phenotype correlations and reduce “noise” in the analysis, although the later does not apply to these data given the unique genotypes of each N2 offspring (Figure 1). Therefore, caution must be applied to directly extrapolating these effect sizes to human genetic studies or even studies in other mammalian systems. Nonetheless, within this CSS-based study, these same factors apply equally to main effects and interaction effects, and the fact that the variance explained was similar between these two classes of QTLs contributes to the growing body of evidence that epistatic interactions play an important role in the genetic architecture of complex traits. This finding has broad implications for the genetic architecture of complex traits, where most studies to date have exclusively identified main effect QTLs. Despite this limitation, clinical applications based on these studies, such as the use of polygenic risk scores, are beginning to be clinically evaluated with some promising early results (Torkamani *et al*. 2018). Our results highlight that these models are likely to remain severely handicapped until they account for interactions, as the individual and cumulative effects of the interacting loci can be equivalent to the main effects, but remain unaccounted for in the models.

While it is relatively easy to lament the absence of epistatic interactions in modeling genetic architecture, the problem is not simply that the models do not account for interactions, but rather that experimentally or statistically identifying such interactions has proven exceedingly difficult. Herein lies the power of this new CSS interaction-based mapping paradigm, which enabled the detection and higher-resolution mapping of otherwise hidden QTLs, revealing the genomic locations of numerous novel loci that interact to control a number of complex traits and the expression of hundreds of genes that were not found using only a marginal analysis strategy. This was all accomplished with a relatively small number of mice for a mapping population, including just ∼150 mice for the trait data and ∼50 mice for the gene expression analyses. Thus, the use of additional CSS crosses and larger mapping populations promises to facilitate the widespread identification of the specific genes and alleles that underlie a broad spectrum of complex traits. These results point to the importance of searching for epistasis in as simplified a genetic context as possible, knowing that even within the limited allelic variation of a single substituted chromosome there is a remarkable genetic complexity contributing to phenotypic variation. As we attempt to move towards more personalized medicine, it will no doubt require a more comprehensive understanding of the many interacting loci throughout the genome, which should be greatly facilitated by this CSS-based mapping paradigm.

## Data availability

RNA-Seq data is available in GEO

Pipeline scripts are available in protocols.io

---

**Supplemental Figure 1. Epistatic interaction between loci on chromosome 6 regulate red cell distribution width in blood**. Red cell distribution width based on genotype at SNP markers (A) rs13478633 and (B) rs13478641 on chromosome 6. Each dot represents one mouse. (C) QTL mapping results for main effects on chromosome 6. The LOD threshold for significance (p < 0.05) was calculated by permutation testing (n=10,000) and is indicated by a horizontal line. Mb position is indicated for both chromosomes 4 and 6 along the x-axis. No significant main effect QTLs were detected. (D) Red cell distribution width based on the combined genotypes at rs13478633 and rs13478641 on chromosome 6. (E) QTL mapping results for interaction effects on chromosome 6. The LOD threshold for significance (p < 0.05)) was calculated by permutation testing (n=10,000) and is indicated by a horizontal line. The most significant interaction QTLs were detected with peaks at rs13478633 (5.2 Mb) and rs13478641 (7.2 Mb). Mb position is indicated for Chromosome 6 along the x-axis. (F) Context-dependent effects of the BB and AB genotypes at rs13478633 and rs13478641. Mean and standard error are shown for each genotype combination. An “A” genotype indicates A/J-derived. A “B” genotype indicates C57BL/6J-derived.

**Supplemental Figure 2. Higher resolution mapping of main effect QTLs regulating the mRNA expression of *Extl1* and *Cadps2***. The main effects for QTLs regulating the expression of *Extl1* to Chromosome 4 and *Cadps2* to Chromosome 6 are denoted by LOD peaks. In our previous study expression of these two genes mapped the chromosome 4 and chromosome 6 in their entirety (Chen *et al*. 2017), but in the current study we were able to map them to a single locus on each of the chromosomes as shown using N2 progeny. (A) The LOD score plot for the *Extl1* QTL is shown, with the LOD score peak located at rs13477965 (126.7 Mb). (B) The LOD score plot for the *Cadps2* QTL is shown, with the LOD score peak located at rs13478695 (32.3 Mb).

**Supplemental Figure 3. Higher resolution mapping of an interaction effect QTL pair regulating the mRNA Expression of *Tmem245***. Using the N2 crosses, we mapped expression of *Tmem245* to interacting loci, one on chromosome 4 and one on chromosome 6. This provided increased mapping resolution in comparison to our previous study where we mapped interacting loci only to the entire chromosomes (Chen *et al*. 2017). The LOD score plot for the *Tmem245* interaction QTL is shown, with the LOD score peaks located at rs13477555 (10.2 Mb) on Chromosome 4 and rs3023840 (147.7 Mb) on Chromosome 6.

**TABLE S3.** TRAIT DATA SUMMARY STATISTICS

| Trait | Female (mean(sd)) | Male (mean(sd)) | T-Test | Wilcoxon Test |
| --- | --- | --- | --- | --- |
| Fasted Weight (g) | 14.09 (0.99) | 16.94 (1.73) | <2.2E-16 | <2.2E-16 |
| Fasting Glucose (mg/dL) | 83.58 (20.39) | 98.79 (24.30) | 6.26E-05 | 6.19E-06 |
| Length (cm) | 88.80 (1.95) | 92.28 (2.73) | 1.28E-15 | 2.04E-14 |
| Fat Pad Weight (g) | 0.043 (0.02) | 0.078 (0.04) | 1.54E-11 | 1.88E-10 |
| White Blood Count (K/uL) | 6.31 (2.20) | 5.95 (2.33) | 3.60E-01 | 3.03E-01 |
| Lymphocytes (K/uL) | 5.15 (1.62) | 4.90 (1.93) | 4.14E-01 | 2.30E-01 |
| Monocytes (K/uL) | 0.31 (0.14) | 0.29 (0.14) | 4.86E-01 | 4.89E-01 |
| Lymphocytes (%) | 83.03 (4.47) | 82.28 (4.14) | 3.20E-01 | 3.36E-01 |
| Basophils (%) | 0.12 (0.13) | 0.17 (0.18) | 5.42E-01 | 5.00E-01 |
| Red Blood Count (M/uL) | 9.32 (1.00) | 9.28 (0.97) | 7.88E-01 | 7.54E-01 |
| Hemoglobin (g/dL) | 14.79 (1.58) | 14.56 (1.73) | 4.30E-01 | 4.83E-01 |
| Hematocrit (%) | 43.63 (4.08) | 44.34 (4.29) | 3.31E-01 | 9.99E-02 |
| Mean Corpus Volume (fL) | 46.88 (1.68) | 47.84 (1.69) | 1.28E-03 | 1.81E-04 |
| Mean Corpuscular Hemoglobin (pg) | 15.85 (0.36) | 15.68 (0.88) | 1.59E-01 | 8.23E-01 |
| Mean Corpuscular Hemoglobin Concentration (g/dL) | 33.87 (1.30) | 32.86 (2.46) | 3.44E-03 | 1.73E-05 |
| Red Cell Distribution Width (%) | 19.11 (1.00) | 19.24 (1.09) | 4.88E-01 | 3.14E-01 |
| Platelets (K/uL) | 562.90 (125.60) | 656.14 (152.13) | 1.73E-04 | 6.25E-06 |
| Mean Platelet Volume (fL) | 5.19 (0.39) | 5.16 (0.44) | 7.06E-01 | 5.19E-01 |
| Granulocyte (K/uL) | 0.75 (0.38) | 0.75 (0.37) | 9.95E-01 | 7.48E-01 |
| Granulocyte (%) | 12.6 (4.06) | 13.17 (3.88) | 4.11E-01 | 1.47E-01 |
| RDWa | 33.62 (1.00) | 34.77 (1.41) | 2.26E-06 | 6.63E-06 |

**TABLE S4.** MAIN EFFECT LOD SCORES FOR THE INTERVALS CONTAINING EPISTATIC TRAIT QTLS

| Trait | Chromosome | SNP | Position (Mb) | LOD | p value |
| --- | --- | --- | --- | --- | --- |
| Lymphocytes (%) | 4 | rs13477644 | 35.28 | 0.393 | 0.182 |
|  | 6 | rs13478739 | 47.03 | 9.535e-5 | 0.982 |
| Red Cell Distribution Width | 6 | rs13478633 | 12.25 | 0.316 | 0.966 |
|  | 6 | rs13478641 | 16.06 | 0.345 | 0.926 |

## Notes

### Competing Interest Statement

The authors have declared no competing interest.

### Summary of Updates

Updated with current submitted draft.

