## Supplementary figures and images for "A novel mapping strategy utilizing mouse chromosome substitution strains identifies multiple epistatic interactions that regulate complex traits"

### Supplemental Figure 1

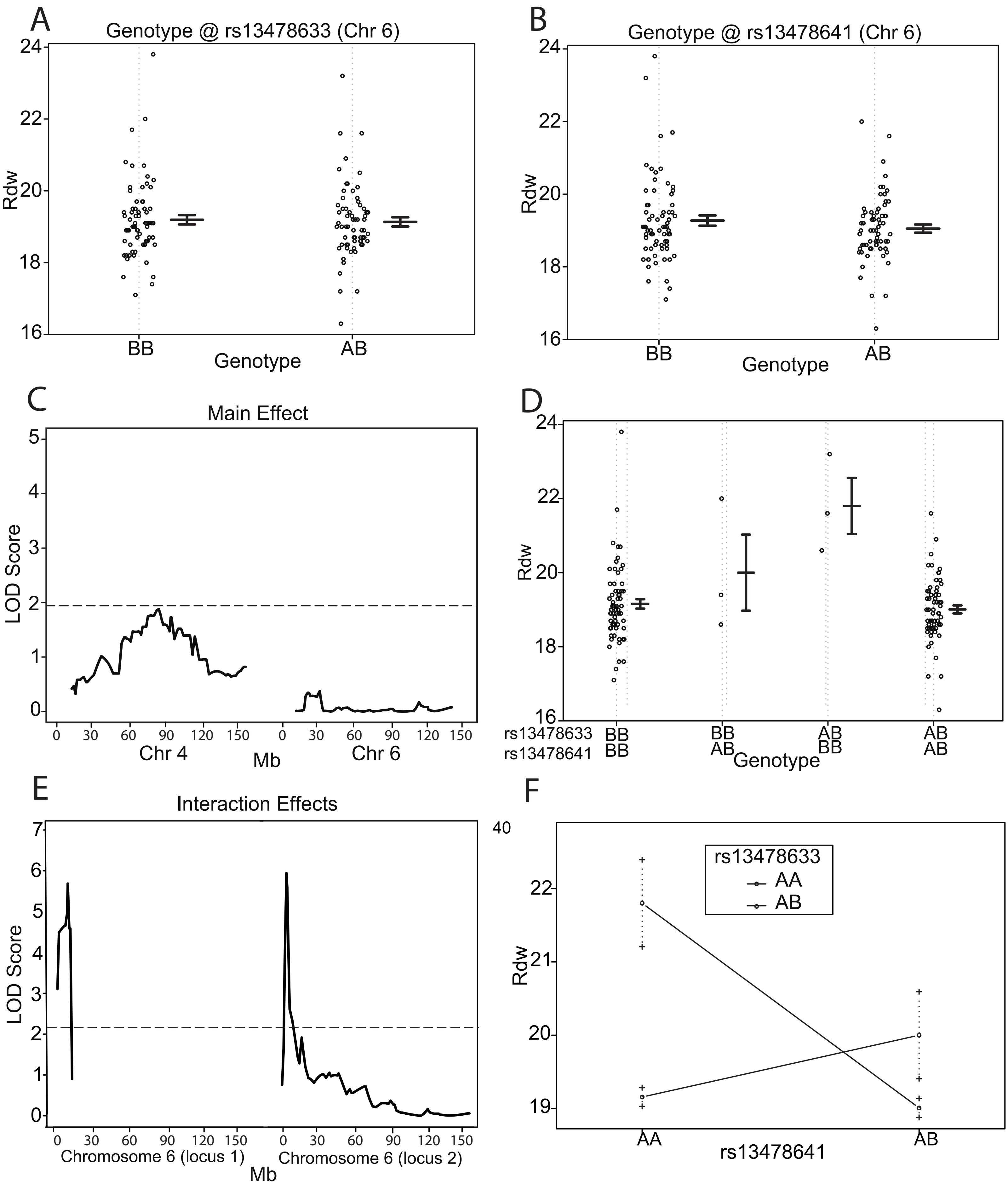

[illegible]

### Supplemental Figure 2

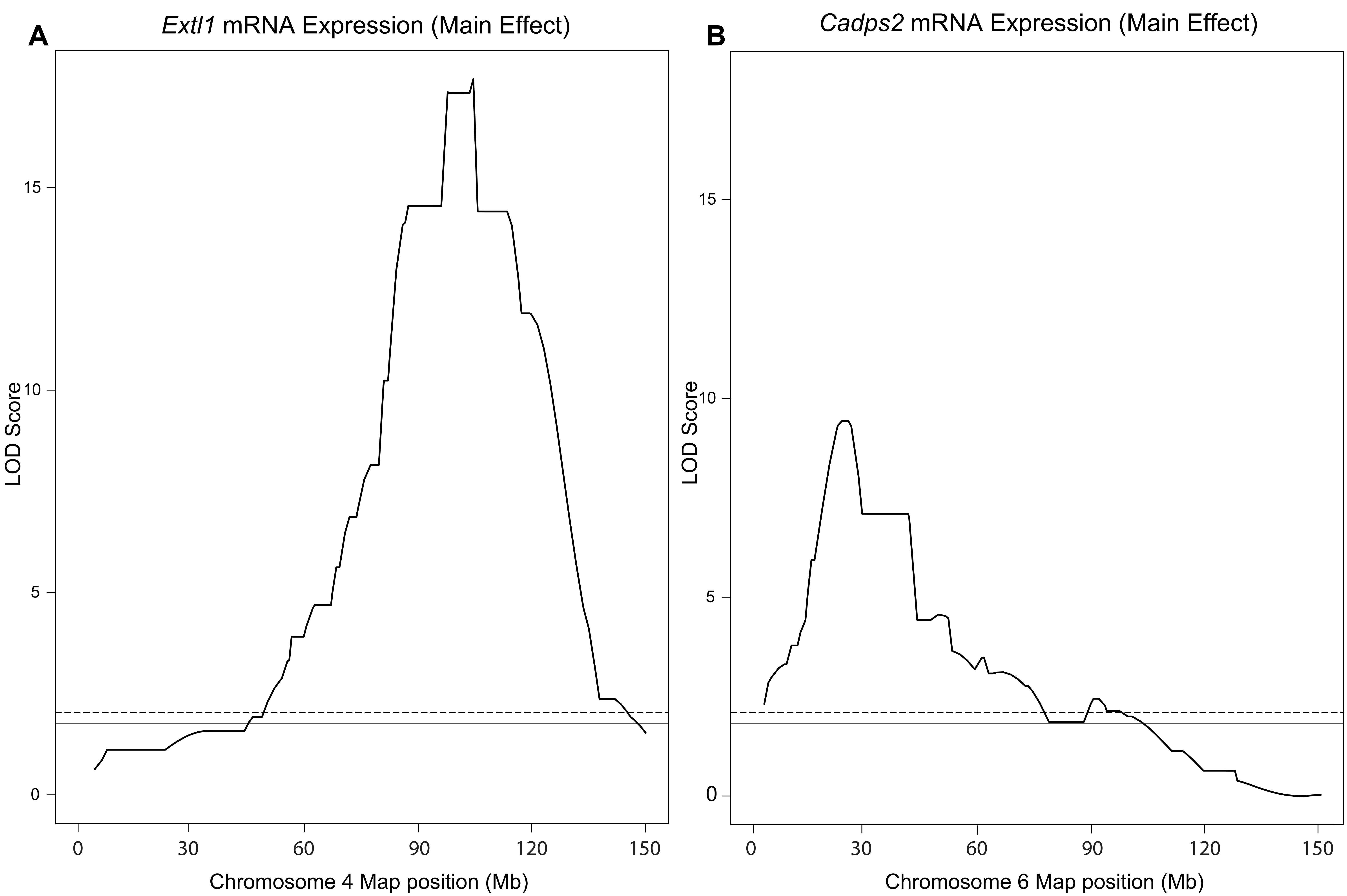

### Supplemental Figure 3

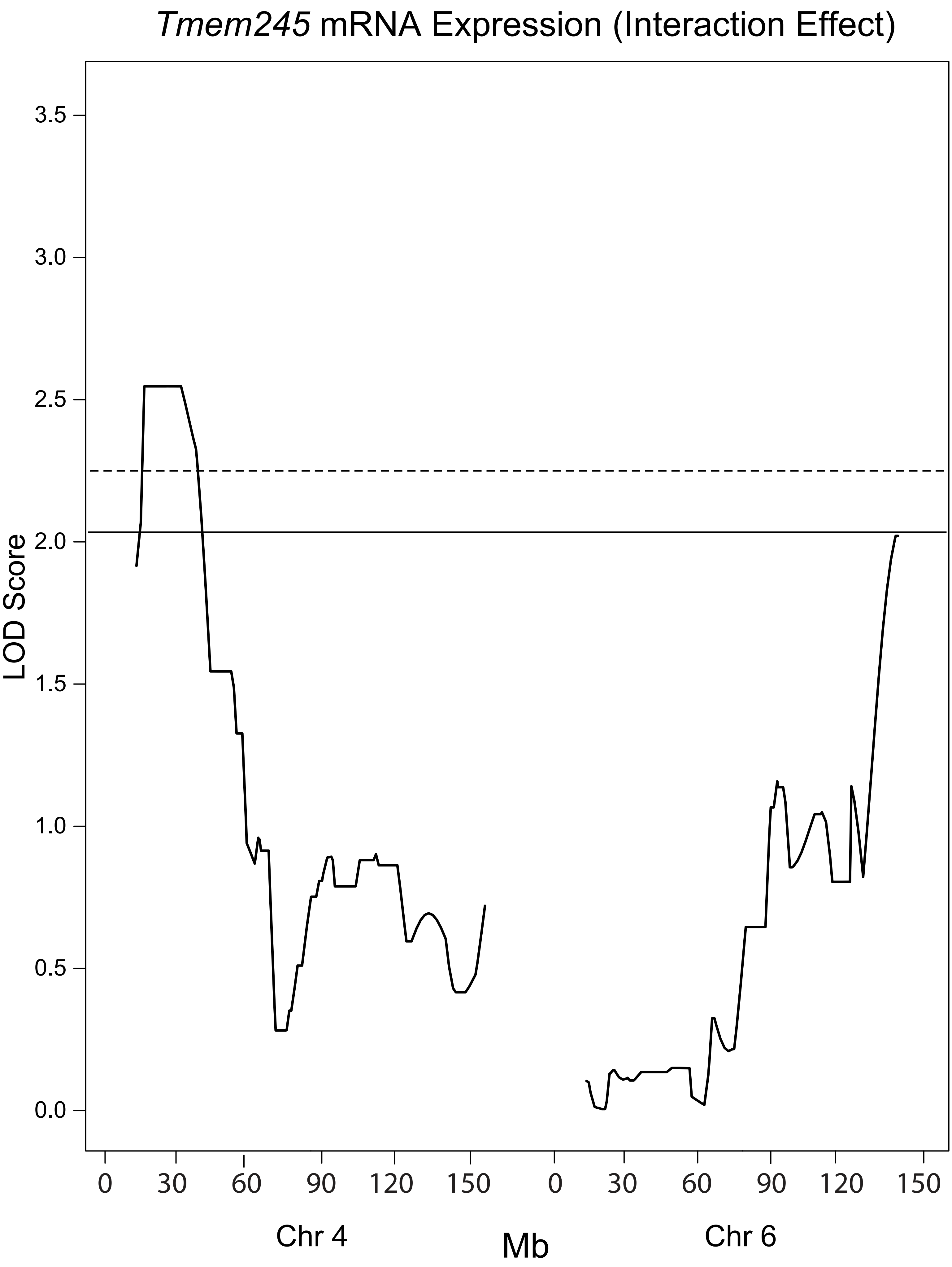
